# Inexpensive and easy method for 6 fragment Golden Gate Assembly of a modular S/MARs mammalian expression vector and its variants

**DOI:** 10.1101/2021.09.04.458594

**Authors:** Adrian Ionut Pascu, Miruna S Stan

**Affiliations:** Department of Genetics, Faculty of Biology, University of Bucharest, Intrarea Portocalelor 1-3, 060101 Bucharest, Romania; Department of Biochemistry and Molecular Biology, Faculty of Biology, University of Bucharest, 91-95 Splaiul Independentei, 050095 Bucharest, Romania

**Keywords:** Golden Gate, Cloning, S/MARS, mammalian expression vector

## Abstract

**Background:** A basic requirement for synthetic biology is the availability of efficient DNA assembly methods. Numerous methods have been previously reported to accomplish this task. One such method has been reported, which allows parallel assembly of multiple DNA fragments in a one-tube reaction, called Golden Gate Assembly. This study aims to further simplify that method and make it more suitable for small labs and students.

**Methods:** Prior to amplification of the parental plasmids used in building the modules were domesticated using a variation of SDM (Site Directed Mutagenesis) called SPRIP. After careful design and amplification of the desired modules, using a high-fidelity polymerase, amplified PCR fragments that enter the one-step-one-pot reaction were stored in Zymo DNA/RNA Shield at -20 degrees C and thawed whenever needed to be used as fragments or modules in the assembly. The fragments were designed to posses unique overhangs using NEB Golden Gate assembly tool and Snapgene, amplification of modules was performed using a Q5 high fidelity polymerase from preexisting plasmids or gene fragments, clean-up of the PCR products (fragments) was performed in one tube per assembly using Zymo DNA Clean and Concentrator-5, assembled using BsaI and T4 ligase, DpnI digestion performed for eliminating the background plasmids that remain after the PCR reaction and the resulting assembled product was transformed into competent *E*.*coli* cells. Transformants were screened using diagnostic digest, transfected into HEK293T cells and the fluorescence was evaluated using fluorescent microscopy and flow cytometry.

**Results:** Herein presented is a simple and inexpensive alternate protocol to build modular plasmids using the Golden Gate Assembly method. A total of p37 S/MARs mammalian expression vectors were designed and constructed using 6 modules previously amplified by PCR and stored in the appropriate buffer to eliminate exo- and endonuclease activity and to protect the DNA from freeze thaw cycles. The existing modules were interchangeable and new modules were easily amplified and stored for use when needed. The mammalian expression vectors constructed showed the desired restriction pattern and GFP expression in bacteria and in mammalian cells. A comparison of 7 pNoname variants was conducted using flow cytometry. Interestingly, no pNoname variant harbouring the SV40 promoter showed expression in tested HEK293T cells. It appears that using the Ef1a promoter in combination with the BGH polyA signal provides the best expression in S/MARS vectors harboring the DTS40 region, as measured by flow cytometry.

**Conclusions:** Provided the design steps are respected and the fragments are stored and labeled appropriately, multiple plasmid variants and combinations of the pre-designed modules can be assembled in one day, easier and using less resources than the established protocols, with good efficiency. The simplicity of the design and the affordability of the method could make modular cloning of plasmid constructs more accessible to small labs and students.

## Background

The principle of Golden Gate Cloning[1] is based on the ability of type IIS enzymes to cleave outside of their recognition site, allowing two DNA fragments flanked by compatible restriction sites to be digested and ligated seamlessly. Since the ligated product of interest does not contain the original type IIS recognition site, it will not be subject to redigestion in a restriction-ligation reaction.

All products that reconstitute the original site will be re-digested, allowing their components to be made available for further ligation, leading to formation of an increasing amount of the desired product with increasing time and cycles of incubation. Since the sequence of the overhangs at the ends of the digested fragments can be chosen to be any 4-nucleotide sequence, multiple compatible DNA fragments can be assembled in a defined linear order in a single restriction-ligation step.

Based on this method multiple standards to construct multi-gene circuits have been developed. Mo-Clo, or the modular cloning standard, is one of these standards [2]. Mo-Clo consists of standard parts, or basic modules, of eukaryotic Promoters, 5’UTR’s, Signal Peptides, CDS’s, 3’UTR’s and Terminator regions, each module containing a standardized 4 nucleotide overhang. Mo-clo or similar standards like Golden Braid 2.0 [3] use 3 levels of successive assemblies.

Level 0, or basic modules (Promoters, 5’UTR’s, Signal Peptides, CDC’s, 3’UTR’s and Terminator regions) are assembled into Level 1 transcriptional units using a Type IIs restriction enzyme. The level 1 transcriptional units can be, in turn, ligated into level 2 multigene constructs using a different type IIs restriction enzyme.

However, although standardized and agreed by the community, these standards may prove too complex for assembling just a few/several fragments or for students that are just getting started in molecular biology or that have a desire for more customization.

In turn, herein described is a simple inexpensive protocol for standardizing and assembling several fragments with good fidelity, in just one day with minimal *hands-on* time.

For this purpose a modular S/MARs mammalian expression vector (called pNoname) has been designed. A S/MARs vector is a type of non-integrating episomal vector that is used for mammalian expression[4][5]. It contains a Scaffold/Matrix Attachment Region which is an element designed to permit the persistence of the vector as an extrachromosomal element within a host cell [6]. It also provides the vector with the ability to remain episomal and be mitotically stable during cell division [5]. Furthermore, these types of vectors can replicate during cell division in certain types of cells. [7].

The modular vector designed in this paper contains 6 fragments or modules that can be easily interchangeable.

In order to facilitate the Golden Gate Assembly, fragments to enter the assembly, previously amplified by PCR can be stored in Zymo DNA/RNA Shield at a ratio of 1:3 (PCR : Zymo Shield) thus eliminating the need to store the fragments in plasmids and facilitating the screening of the desired mutants.

Zymo DNA/RNA shield protects DNA from freeze thaw cycles, stabilizes nucleic acids and deactivates nucleases. [8]. DNA overhangs of several bases designed in primers can further protect the PCR products from exonucleases thus leaving the type IIs restriction sytes intact.

After designing the primers to possess specific 4 base pair overhangs and accommodating the DNA fragments that will enter the assembly reaction to lack any other specific type IIs restriction sites (depending on the type IIs restriction enzyme used) [9] the fragments can be cleaned, assembled and the resulting product transformed into competent *E*.*coli* cells (*home-made* or purchased) to generate the desired mutants.

Multiple plasmids can be constructed at the same time provided that certain rules are followed and provided that the correct fragments are mixed for each reaction.

Fragment assemblies (called F or modules) and transformations can be performed in the same day. The core fragment which contains the bacterial origin of replication and the antibiotic gene can be amplified by PCR. Multiple types of core fragments could be used, with interchangeable antibiotic resistance genes. Of the amplified PCR products stored in ZS (Zymo Shield), 20 µl can be added to the assembly reaction (of each fragment type). Before assembly, the PCR mix (mixed fragments) is cleaned and concentrated and 20 µl of eluate is used in the GG assembly reaction. Multiple reactions can be made at the same time. After the assembly, DpnI digestion is performed to remove the background plasmid that was used as a PCR template and then 2-5 µl is transformed into competent *E*.*coli*.

## Methods

### pNoname design and construction

In order to provide a backbone for future experiments a modular S/MARs vector called pNoname was designed. All *in silico* designs were made using the Snapgene software [10]. The vector consists of 6 modules, designed to be easily interchangeable in a short amount of time and with minimum financial effort.

1. One module consists of the required bacterial origin of replication for replication and maintenance in bacteria and the antibiotic resistance gene for selection.
2. One module consists of the promoter and enhancer region.
3. One module consists of the reporter gene (cGFP) with a Kozack consensus sequence.[11]
4. One module consists of a shortened Nuclear Matrix Attachment Region, MARS5 [7] directly downstream of the gene.
5. One module consists of the Terminator and the poly adenylation sequence.
6. One module consists of the SV40 (Simian Virus 40) DTS (DNA nuclear targeting sequence), a 72 bp repeat of the early enhancer, that has been shown to mediate nuclear transport of the inserted DNA. [12]

The primers for amplifying the fragments or modules were designed using the New England Biolabs Golden Gate Assembly Tool [13] in order to generate specific 4 base pairs overhangs and were further modified to accommodate other requirements (GC content, primer TM etc) using Snapgene. The standardized 4 base pairs overhangs are highlighted in (Figure 1). After digestion with BsaI and ligation with T4 ligase the BsaI recognition sequence is eliminated from the final construct thus preventing redigestion. The primers used for the amplification of the modules are shown in Supplementary Materials [Additional File 3].

**Figure. 1.**
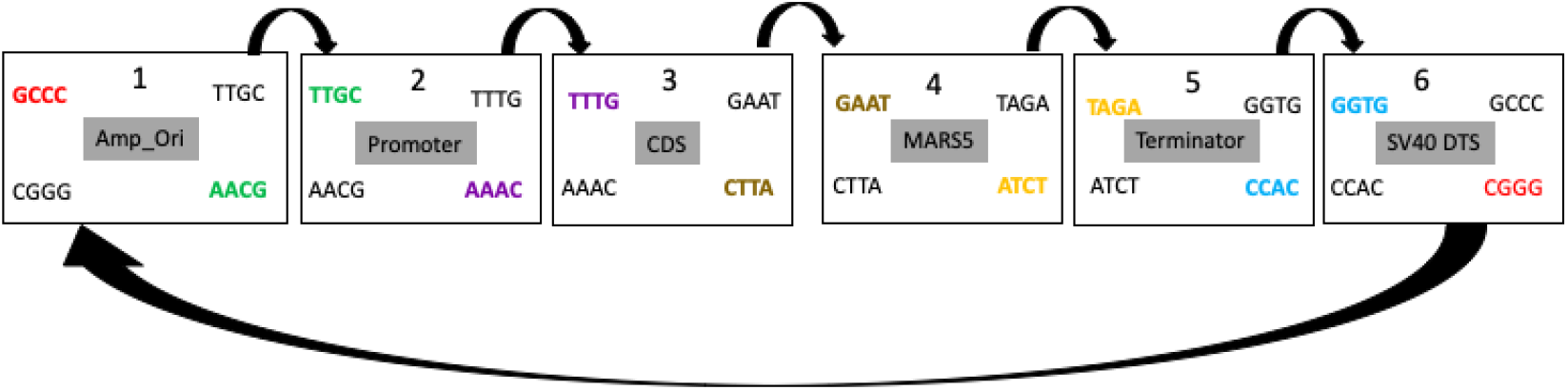
Overhangs generated for each module with the assemble direction given in arrows. Overhangs are given in colours. Each colour is specific (yellow binds to yellow, red binds to red etc)The design includes 6 modules that can be assembled (Amp_ori, Promoter, Coding sequence, S/MARS region, terminator or poly adenilation sequence and the DTS40). Fragments can be easily substituted for new ones using the 5’ “tails” of the primers”

A schematic representation of pNoname variant 1 (pNoname_CMV_GFP_BGH) is shown in Figure 2.

**Figure 2.**
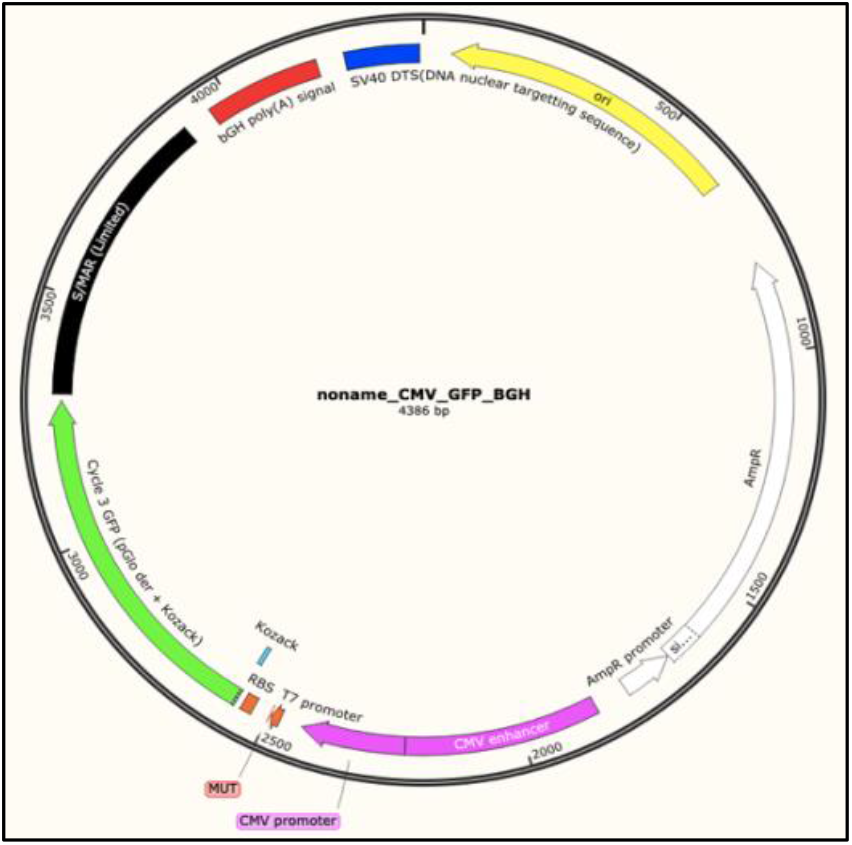
pNoname variant CMV_GFP_BGH after assembly that contains all 6 modules (Promoter region, Coding seqence, MARS5, Terminator and poly adenilation signal, SV40 DTS and the core fragment (bacterial origin of replication and antibiotic resistance gene)

In total, 7 pNoname variants were constructed with the GFP gene under the control of one combination of Promoter and Terminator (polyadenylation sequence) as follows.

3 pNoname variants were under the control of the EF1A promoter and SV40, BGA and BGH poly-A sequences respectively, 3 pNoname variants were under the control of the SV40 promoter and SV40, BGA and BHG poly-A sequences respectively and one pNoname variant under the control of the CMV promoter and enhancer and BGH poly-A sequence.

### Domestication (removal of internal BsaI sites)

In order to enter the Golden Gate Assembly, all other BsaI sites except the ones containing the overhangs must be eliminated from the fragments.

Domestication of the plasmids was performed using primers FPCTokillmyBsaI, RPCTokillmyBsaI, FTokillBsaI, RtokillBsaI (provided in [Additional file 3]) with a modified version of Site Directed Mutagenesis, called SPRIP (Single Primer Reaction in Parallel) [14]. Two plasmids were domesticated, pUC18 and pcDNA3.1(+) (ThermoFischer Cat.No V79020). The primers were designed to be complementary to each other and to contain a single base pair mutation that will eliminate the BsaI from the ampicilin resistance gene of the pUC18 plasmid and eliminate the BsaI site just downstream of the T7 promoter of pcDNA3.1(+) respectively. Single primer PCR was performed in 2 different tubes for each plasmid using Q5 2X Master Mix (New England Biolabs Cat.No M0492S). After the amplification, the PCR tubes contents were mixed and allow to re-anneal using the following cycle (5 min at 95° C; 1 min at 90° C; 1 min at 80°C, 0.5 min at 70°C, 0.5 min at 60°C, 0.5 min at 50°C, 0.5 min at 40°C, hold at 37°C). After the re-annealing step, DpnI (New England Biolabs Cat.No R0176S) digestion was performed prior to transformation to eliminate the parental plasmid. Transformed cells (New England Biolabs *E*.*coli* Turbo Cat.No C2984H) made competent using the protocol provided in [Additional file 1] were placed in the incubator overnight at 37° and colonies were screened for desired mutants using BsaI-HF®v2 (New England Biolabs Cat.No R3733S).

### Construction of modules (fragments) for assembly

The Amp_Ori region was amplified from pUC18 after it’s *domestication* (removal of any internal BsaI sites). The other modules used in this experiment, CMV promoter and Enhancer, the SV40 promoter, the SV40-DTS, the BGH terminator region and the SV40 terminator region were amplified from pcDNA3.1(+) after domestication. The BGA polyadenylation signal was amplified from plasmid C7 (MXS Chaining Kit; Addgene reference #62424)[15] and the Ef1A promoter was amplified from plasmid C4 (MXS Chaining Kit; Addgene reference #62421).

Gfp_Kozack (The reporter gene coding for a green fluorescent protein) and MARS5 were synthetized by Twist Biosciences. Primers were used to further multiply the sequences from pNoname.

All amplifications were made using a high-fidelity polymerase (Q5 2X Master Mix, New England Biolabs) according to the manufacturers’ recommendations in a 25 µl final reaction volume.

All primers used in this study are provided in [Additional File 3].

### Construction of pNoname variants

After amplification, 20 µl of the PCR product that contained the specific modules were stored in Zymo DNA/RNA shield (Zymo Research Cat.No R1100-50) in the freezer at -20 °C, at a ratio of 1:3 (DNA : Zymo Shield / 20 µl DNA:60 µl Zymo Shield). O volume of 5µl of the PCR product was loaded on a gel for confirmation of the reaction. 20 µl of each module (PCR product + Zymo Shield) (6 in total) were mixed in a single tube and cleaned using the Zymo Research DNA Clean and Concentrator -5 kit (Cat.No D4013). Each tube contained fragments from a single pNoname variant. A total of 7 pNoname variants were constructed. (pNoname_CMV_GFP_BGH, pNoname_SV40_GFP_SV40, pNoname_SV40_GFP_BGH, pNoname_SV40_GFP_BGA, pNoname_Ef1a_GFP_BGH, pNoname_Ef1A_GFP_BGA, pNoname_Ef1A_GFP_SV40). The assembly reaction was set up as follows: 20 µl of mixed purified fragments (6 fragments in total), 1.5 µl (600 U) of T4 ligase (New England Biolabs Cat.No M0202S), 1 µl (20 U) of BsaI-HF®v2, 2,5 µl of T4 Ligase Buffer. 5 µl of purified fragments were kept for gel agarose confirmation. The assembly cycle was: (1 min at 16° C + 1 min at 37°C) X 30 cycles followed by a 5 minute soak time at 60°C. DpnI digestion was performed to eliminate any background plasmids from the PCR reaction using 1 µl of DpnI (37°C for 15 minutes followed by 5 minutes at 60°C) and transformed with 5µl of assembly product using *E*.*coli* Neb Turbo made competent by the transformation protocol in Supplemental Materials. Cells were plated on antibiotic selection plates with LB Agar containing 100 µg/ml ampicilin and left in the incubator overnight at 37°C.

Mutants were confirmed by restriction digest on a 1% agarose gel and by amplifying the regions using primers provided in Table 1.

**Table 1 :**
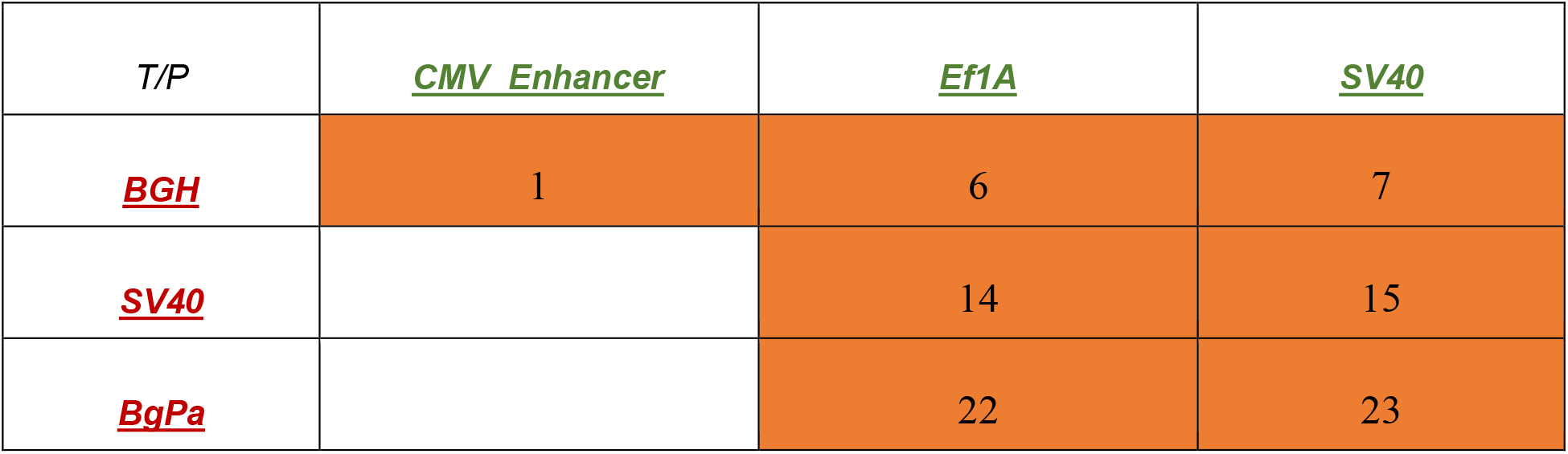
Variants constructed (orange background) : pNoname variant 1 (CMV promo/enhancer + BGH polyA), 6, 14 and 22 (Ef1A promoter and BGH terminator, SV40 terminator and BgpA terminator respectively) and 7, 15, 23 (SV40 promoter and BGH terminator, SV40 terminator and BgpA terminator respectively).

| *T/P* | *CMV Enhancer* | *Ef1A* | *SV40* |
| --- | --- | --- | --- |
| *BGH* | 1 | 6 | 7 |
| *SV40* |  | 14 | 15 |
| *BgPa* |  | 22 | 23 |

The detailed design and assembly protocol is provided in [Additional file 2].

The variants constructed are presented in Table 1.

### Fluorescence testing for pNoname variants

Fluorescence was determined visually in E.coli BL21(DE3) for the variant harboring the CMV promoter and enhancer by induction with 100mM IPTG.

HEK293T cells were cultured in a humidified atmosphere with 5% CO_2_ at 37 °C, using complete Dulbecco’s Modified Eagle Medium (DMEM; Gibco/Invitrogen, Carlsbad, CA, USA) supplemented with 10% fetal bovine serum (Gibco/Invitrogen, Carlsbad, CA, USA). The growth medium was replaced with a fresh one every 2 days until 80% confluence was reached. Then, a 0.25% (w/v) Trypsin 0.53 mM EDTA solution (Sigma-Aldrich, St. Louis, MO, USA) was used to detach and split the cells for future sub-cultivations. Cells were cultured at a density of 105 cells/cm2 in 6-well plates and left to adhere overnight. After 24 h cells were transfected with 2 µg of each plasmid per well using Jet Optimus (Polyplus Transfection cat. no. 101000051) according to the manufacturers’ recommendations. The green fluorescence was visualized with an inverted fluorescence microscope Olympus IX71 (Olympus, Tokyo, Japan) at 27 h, 48 h, 147 h and 243 h post-transfection. This long period of evaluation was possible by passaging the cells using a 10-times dilution after each time point. Also, the median and number of fluorescent cells for each variant was measured. using a 20-times dilution of cells harvested after each time interval using the Accuri C6 (Beckton Dickenson) flow cytometer. Measurements were done in triplicate and the median FITC was the media of the 3 measurements for each variant.

## Results

### Construction of modules (fragments) for assembly

The fragments (modules) needed for assembly were amplified using a high-fidelity polymerase in a final reaction volume of 25 µl, according to the suppliers’ recommendations. 5 µl of each reaction was loaded onto a 1% agarose gel next do a DNA ladder to confirm successful amplification. 20 µl of each reaction was stored at a ratio of 1:3 in Zymo DNA/RNA Shield and aftert proper labelling, the tubes were placed in the freezer at -20 °.

### Construction of pNoname variants

The 7 pNoname variants were constructed using 20µl of each module previously amplified and stored in Zymo Shield. The six modules were loaded in each of the 8 tubes corresponding to each variant to assemble. A gel electrophoresis pre and post assembly confirmed the successful Aassembly of the variants (Figure 5). Data not shown for pNoname_CMV_GFP_SV40, pNoname_CMV_GFP_BGH, pNoname_Ef1a_GFP_Sv40, pNoname_Ef1a_GFP_BGA, pNoname_Ef1a_GFP_BGH,

**Figure 3.**
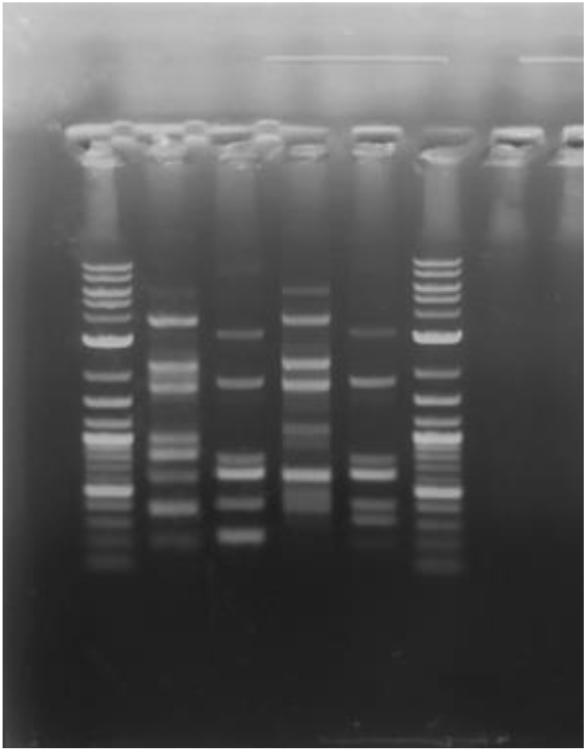
Lane 1 Ladder 1 kb plus New England Biolabs, Lane 2 pNoname_SV40_GFP_SV40 post assembly, Lane 3 pNoname_SV40_GFP_SV40 pre assembly, Lane 4 pNoname_SV40_GFP_BGH post assembly, pNoname_SV40_GFP_BGH pre assembly

**Figure 4.**
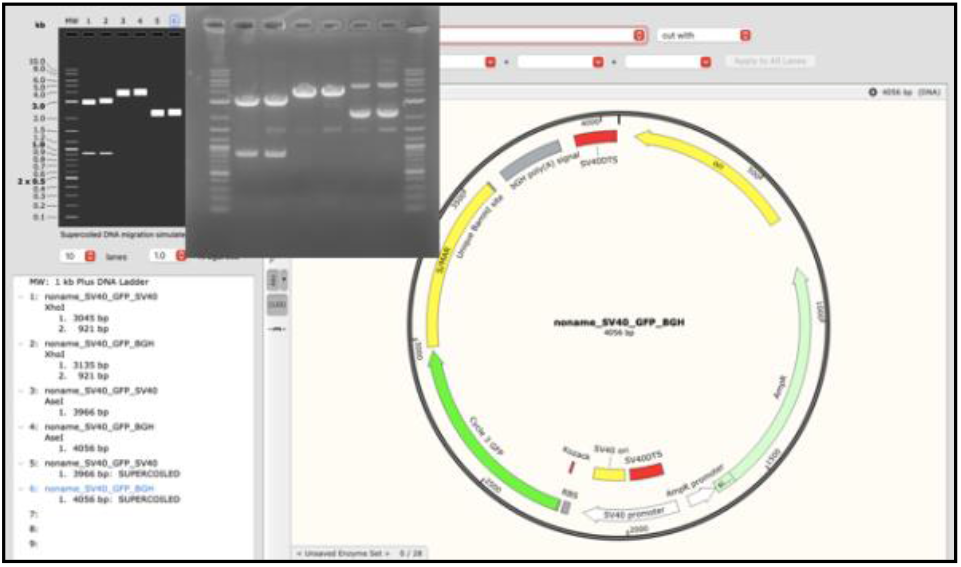
pNoname variants. Lane 1 Ladder 1kb plus New England Biolabs, Lane 2 pNoname_SV40_GFP_SV40/XhoI, Lane 3 pNoname_SV40_GFP_BGH/ XhoI, Lane 4 pNoname_SV40_GFP_SV40/ AseI, Lane 5 pNoname_SV40_GFP_BGH/AseI, Lane 6 pNoname_SV40_GFP_SV40 non-digested, Lane 7 pNoname_SV40_GFP_BGH non-digested, Lane 8 Ladder 1kb plus New England Biolabs

**Figure 5.**
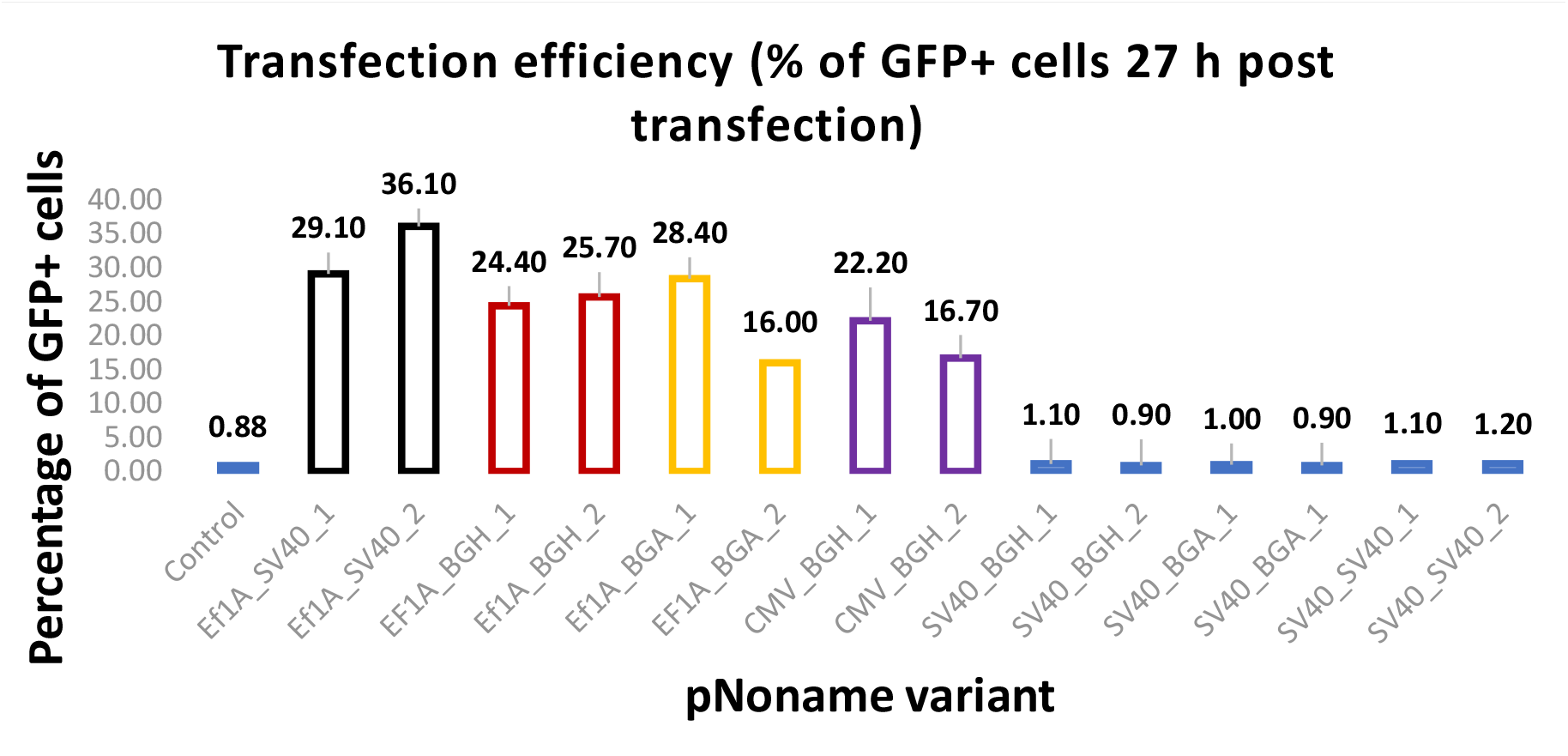
% + cells 27h post transfection for measured pNoname variants. The X axis represent the pNoname variants and the Y axis represents the percentage of positive cells. Expperiments were performed in duplicate. Control is the background fluorescnece (non transfected cells)

After transformation 2-4 colonies from each of the four plates were harvested for screening. Desired mutants were confirmed by restriction digest according to their individual DNA sequences.

### pNoname variants fluorescent expression testing

The pNoname variant CMV_GFP_BGH (pNoname_1) was designed to contain a T7 promoter for ease of screening in bacteria. Fluoresecence was determined visually and by flow cytometry on the other variants as described in Materials and Methods.

Fluorescence was measured in duplicate at T0 (27 h post transfection) for each variant. At T0 the percentage of transfected cells were 32% for pNoname_14 (Ef1A_GFP_SV40), 25 % for pNoname_6 (Ef1A_GFP_BGH), 22% for pNoname_22 (Ef1A_GFP_BGA) and 20% for pNoname_1 (CMV_GFP_BGH).

Interestingly, for the variants under the control of the SV40 promoter no GFP cells were observed in HEK293T compared to the background fluorescence (Figure 5). This could be due to the fact that the pNoname variants, through the DTS40 region, contain an additional region of the SV40 promoter, promoter shown to have a somehow weak expression in HEK293T cell [16]. The large T antigen expressed by the HEK293T cells [17] could also present an issue regarding the poor expression in this article, however this remains to be determined.

Median fluorescence intensity was also measured at the specified timelines for pNoname variants 6,14,22,1. No further measurements were performed for the SV40 promoter variants (7,15,23) following the poor expression at 27 hours.

At 27 hours post-transfection, the best expression was provided by pNoname EF1A_SV40 with a median FITC value of 50000 followed by Ef1A_BGH (median FITC of 37000), Ef1A_BGA (median FITC of 32000) and lastly, CMV_BGH (median FITC of 19000).

At 48 hours post-transfection, the median FITC of Ef1A_SV40 started to decline, the Ef1A_BGH and Ef1A_BGA peaked at approx. 100000 and 76000 median FITC respectively and CMV_BGH remained constant. At T3 (147 hours) the expression for GFP in all 4 variants declined followed by a further decline at T4 (243 hours). Data is shown in (Figure 5 and Figure 6). Fluorescence microscopy of the variants also seem to confirm these findings (Figure 7).

**Figure 6.**
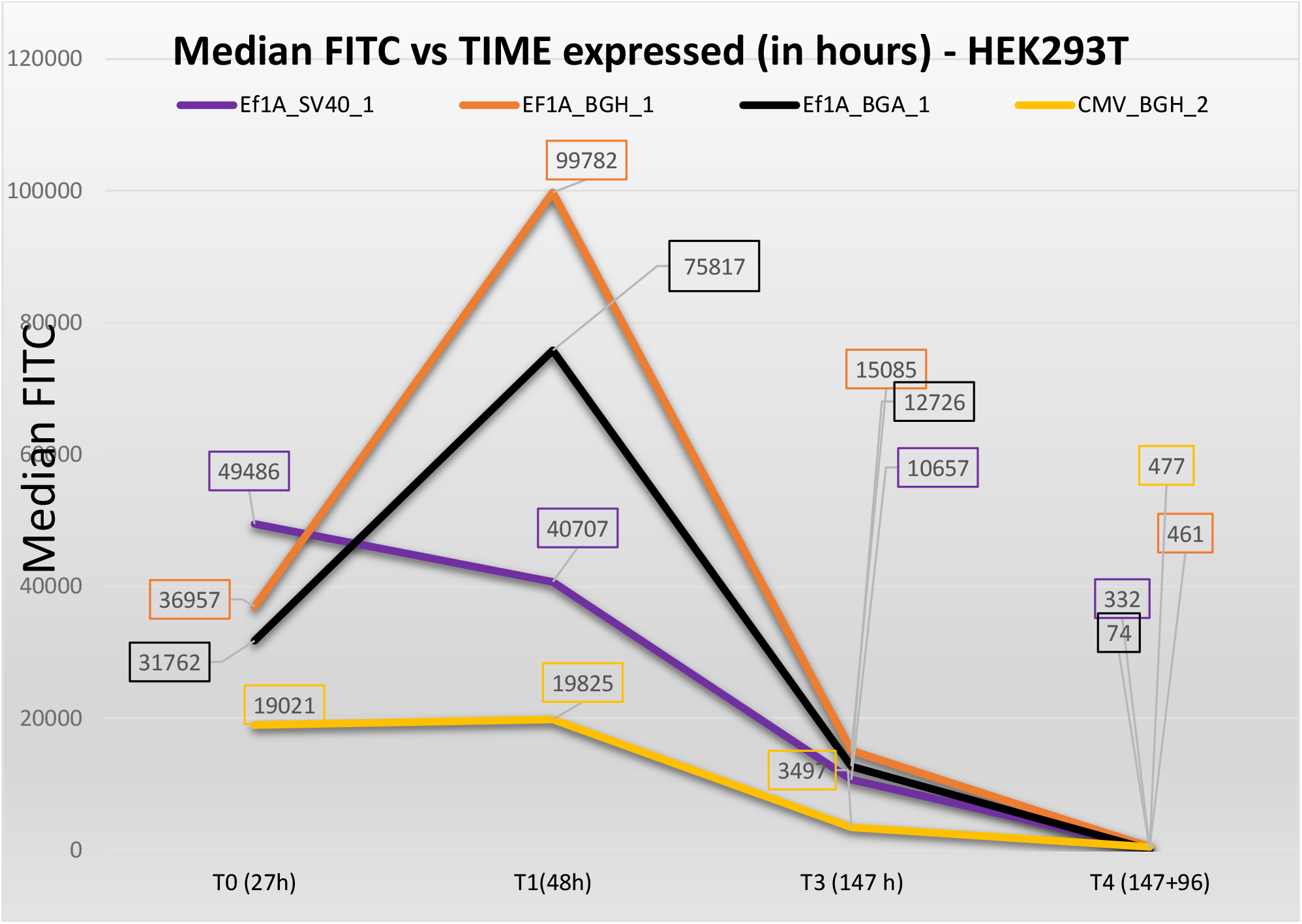
Median FITC as measured by Flow Cytometry. Ef1A_SV40 showed best expression after 27 hours. However, expression for EF1A_SV40 started to decline at 48 h post transfection. Ef1A_BGH showed a peak at aprox 10000 median FITC at the 48 h mark, followed by Ef1A_BGA and Ef1A_Sv40. Although constant CMV_BGH had the weakest expression in the tested conditions at both 27 h and 48 h.

**Figure 7.**
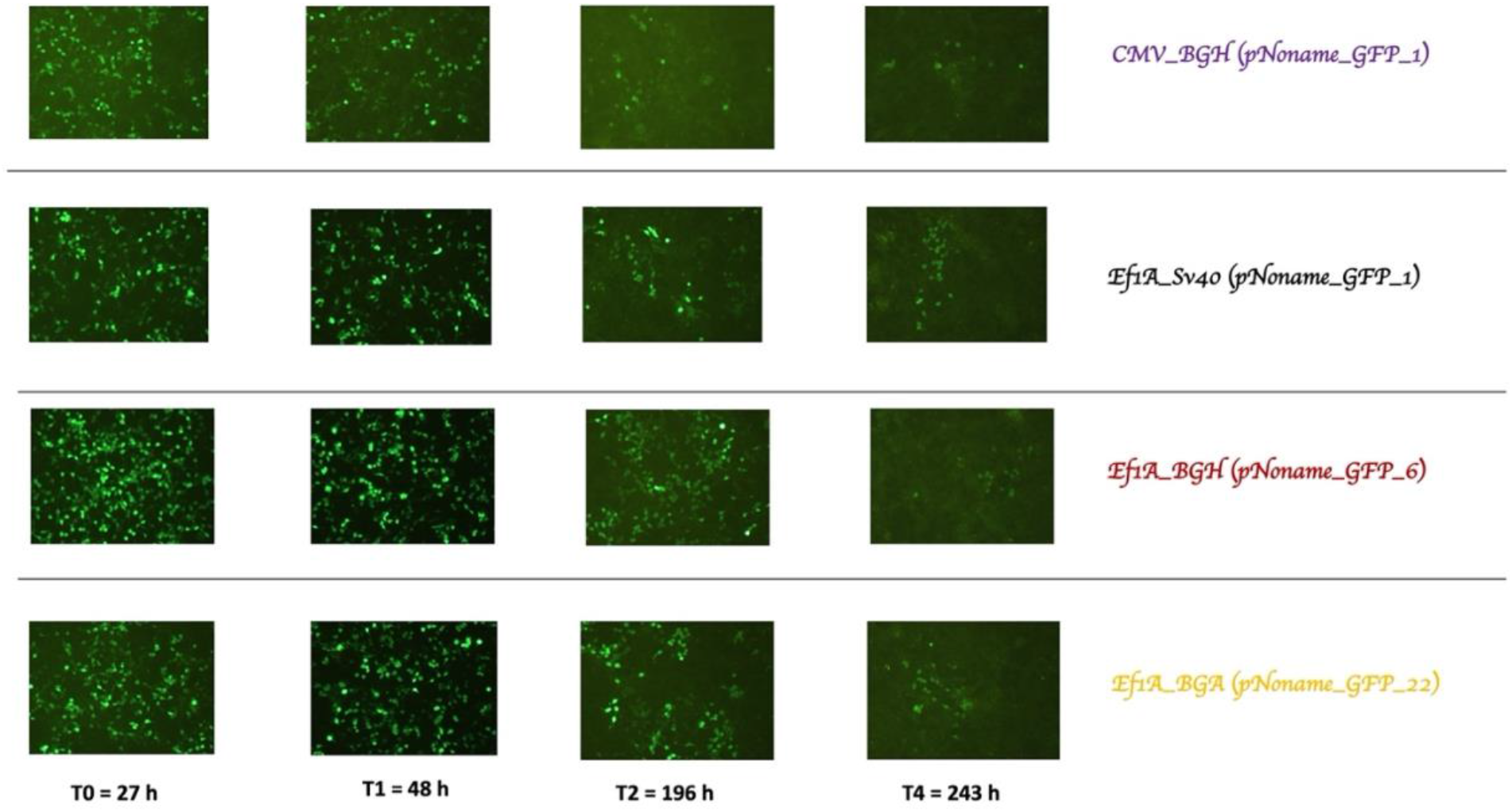
Fluorescence microscopy for the 4 plasmids transfected at 27 h, 48 h, 147 h, and 243 h in Hek293T. First row : CMV_BGH, second row EF1A_SV40, third row EF1A_BGH, forth row EF1A_BGA. Microscopy seems to confirm the findings regarding fluorescence expression from flow cytometry.

## Discussion

The need for cheap, easy methods to design and build various expression vectors is still present in molecular and synthetic biology, especially for small laboratories that lack the capacity to implement large scale screening and/or construction of many vectors at a time and complicated gene circuits. By careful design and with cost savings in mind, small laboratories and students can build vectors without the need for multiple restriction enzymes, laborious screening and using just a few selected products.

The protocol presented here could be such an alternative. Counterintuitively, by building the plasmids from the ground up every time instead of cloning a single gene, using stored modules amplified by PCR (including the core element consisting of the bacterial origin of replication and the antibiotic resistance gene), instead of using the concept of vectors and inserts, the workflow could be simplified.

The design could be adapted easily to accommodate more/less modules. Using just two primers designed to possess the right overhangs one or more fragment types could be eliminated from the final construct with ease.

The fact that equimolar amounts of both fragments and „vector backbone” could be introduced into the assembly reaction and the amount of DNA could be approximated by gel electrophoresis alone, would eliminate the use of a spectrophotometer.

Storing the amplified PCR fragments or modules in an environment that is endo-and exonuclease free, protecting the nucleic acids from freeze thaw cycles, amplifying further modules with a high fidelity polymerase that has proof reading capabilities when needed using just 1µl of parental plasmid, cleaning the PCR products just one time per assembly reaction, customizing the modules through careful design and using just one restriction enzyme for molecular cloning could be a reliable and cheap method for multiple modular vector construction in a very short time, reducing hand-on time and costs.

## Conclusions

This method demonstrates that modular expression vectors could be constructed easily, affordably and with minimal *hands-on* time by even the most under funded and under equipped laboratories. By carefully optimizing the design and methods used, multiple modular expression vectors with interchangeable parts can be build by anyone using minimum equipment and reagents. This can be further used for optimizing gene expression in different cell lines at a very low cost.

