## Additional File 1 for "Inexpensive and easy method for 6 fragment Golden Gate Assembly of a modular S/MARs mammalian expression vector and its variants"

**Day 1**

1.Prepare fresh bacteria (culture or plate)

**Day 2**

2. Inoculate new culture from O/N culture or plate
3. Incubate new culture at 37C with shaking until it reaches OD600 = 0,3-0,4 (3-4 h)
4. Load 2ml/1,5ml of new culture into 2 ml tube
5. Put an extra 2 ml/1,5 ml tube, DNA, and 100ul of Buffer on ice.
6. Ice everything for 10 minutes
7. Pellet cells in centrifuge (6 minutes at 1000G 4 degrees Celsius)
8. Discard supernatant
9. Resuspend pellet gently in 100ul of chilled BUFFER
10. Divide culture into two tubes, label + and –

***Note. To increase the transformation efficiency cells with CaCl2 can be incubated overnight PRIOR to adding DNA***

11. Add 2ul DNA (100 pg-100ng) to the + tube
12. Mix gently by flicking tube
13. Incubate on ice for 30 minute
14. Heatshock for 30 sec at 42 C (time varies depending on the strain used) *E.Coli* Turbo – 30 sec, *E.coli* Bl21 10 sec
15. Leave on ice for 5 minutes

16. Add 950ul room temp LB(LB24) to both + and -
17. Place in incubator at 37C for 1 hour with shaking
18. Divide plate into a 1⁄4 region and a 3⁄4 region with sharpy on bottom
21. Label the 1⁄4 region – and the 3⁄4 region +
22. Plate 75ul of the + tube on the 3⁄4 region and 25ul of the – tube on the 1⁄4 region

23. Incubate overnight at 37C on selective media.

**Tips:**

1. Be **VERY GENTLE** after adding BUFFER they are sensitive to mechanical stress now.

2. **Perform ALL steps (5-15 on ICE).**

3. At step 4 depending on the expected no’ of colonies dilute the outgrowth 50 ul + 950 ul LB and plate 75 ul to + from the dilution.

**Buffer :**

Put 0,72 g Cacl2 in 50 ml LB . Filter sterilize or autoclave.

🡺 take 8,5 ml and add to it 1,5 ml Glycerol 🡺 ~ 100 mM Cacl2 15% Gycerol.
