## Additional File 2 for "Inexpensive and easy method for 6 fragment Golden Gate Assembly of a modular S/MARs mammalian expression vector and its variants"

**Six fragment Golden Gate Assembly Protocol**

- **Step 1 – Design primers and modules according to Golden Gate Rules**. (Eg. Domesticate the regions that go into the reaction, design the overhangs etc).

1) Neb Golden Gate Assembly tool can be used for Golden Gate Assembly primer design (<https://goldengate.neb.com/#!/>) [1]

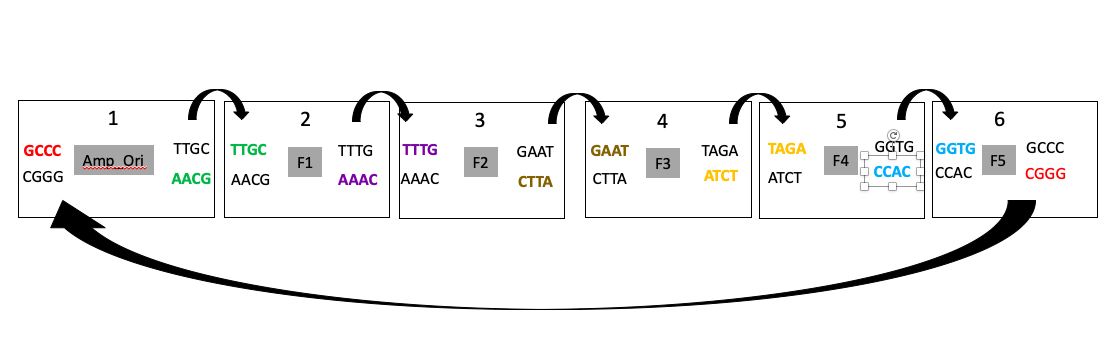

Figure 1 Golden Gate Assembly 6 fragment design. **In bold and coloured : Overhangs**

Design primers in the following way :

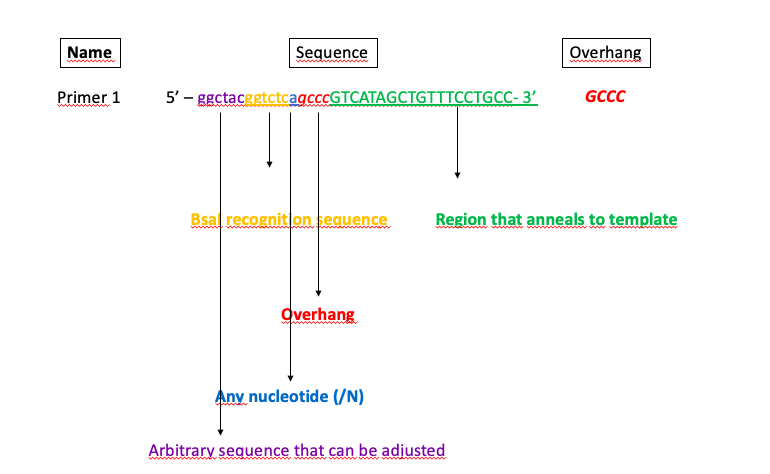

2) For plasmid domestication (removal of any type II S restriction enzymes that are used, in this case BsaI, Site directed mutagenesis [2][3] or SPRIP [4] can be used.
3) Primers used in this article (Supplemental information Primers used) may be adjusted and overhangs can be kept provided TM is taken into consideration and the annealing part of the primer anneals perfectly to the template.

Example :

AbR_Ori fragment forward 🡺 5’ ggctac**ggtctca***gccc*ctaactCCAGCAAAAGGCCAGGAACC 3’

AbR_Ori fragment reverse 🡺 5’ ggctac**ggtctct***gcaa*GTCGACGTCAGGTGGCAC 3’

Promoter fragment forward 🡺 5’ ggctac**ggtctct*ttgc***TTCGCGATGTACGGG 3’

Promoter fragment reverse 🡺5’ ggctac**ggtctca*caaa***attatttcAAGTTTAAACGCTAGCCAGCTTG 3’

**In red** = part to replace

- **Step 2 – Amplify the modules (regions) from plasmids or/and genes**

1) Amplify the regions or fragments using a High Fidelity Polymerase (Eg. Neb Q5 2X Master Mix cat. No M0492S) into a 25 µl reaction.
2) Use 5 µl of the reaction to confirm the amplification on a 1% agarose Gel
3) Mix the remaining PCR product (20 µl) with Zymo Shield (cat. No. R1100-50) at a ratio of 1 PCR : 3 Zymo Shield. Eg 20 µl PCR product : 60 µl Zymo Shield).
4) Store in the freezer and use as module for Golden Gate Assembly.

- **Step 3 – Golden Gate Assembly**

1) Take your fragments/modules out of the freezer.
2) Mix 20 µl of each fragment (including the fragment that contains the origin of replication and antibiotic resitance gene) into a single tube (multiple tubes can be used for multiple reactions)
3) Clean and concentrate the mix using Zymo DNA Clean and concentrator – 5 (Cat no. D4013)
4) Elute the mixed cleaned fragments in 25 µl of water or Elution Buffer
5) Keep 5 µl for Gel Electrophoresis
6) Add into a PCR tube the reaction mix

🡺 1 µl BsaI Hf V2 (Neb cat No. R3733S) , aprox. 20 U

🡺 1,5 µl T4 DNA Ligase (Neb cat No. M0202S), aprox 600 U

🡺 2,5 µl T4 DNA Ligase Buffer

🡺 20 µl Purified Fragments (inluding the AbR_Ori fragments which will act as a backbone).

GGA (Golden Gate Assembly Reaction Conditions)

1 min at 37 degrees C

30 cycles

1 minute at 16 degrees C

5 minutes at 60 degrees C soak time

7) Take 5 µl of the assembly reaction and compare to the non-assembled fragments on a 1% agarose gel
8) Put the remaining 20 µl of assembly reaction in the fridge overnight or proceed to the next step

- **DpnI digestion**

1) Add 1 µl DpnI (Neb Cat No. R0176S) to the assembly
2) Incubate at 37 degrees C for 15 minutes to remove any background plasmid from your PCR amplifications
3) Repeat the soak time (5 minutes at 60 degrees C)

- **Step 4 – Transformation**

1) Use 2-5 µl to transform competent *E.coli* cells
2) Pick single colonies and grow overnight
3) Screen for mutants using diagnostic restriction digest.

***Note :***

- For a cheaper version competent cells can be made in house using Transformation Protocol 1
- I’ve only tried this approach using 6 fragments (modules). My original Golden Gate Design is based on this.
- Expect a reduced number of collonies if using Transformation protocol 1. I usually screened 1-4 colonies for plasmids I’ve build. I always got the desired mutants according to diagnostic digest. I didn’t sequence any of my plasmids.
